# Type 1 Regulatory T cells induced by intestinal epithelial cells respond to food antigen

**DOI:** 10.1101/2025.07.23.666039

**Authors:** Tingyue Zhou, Guorong Zhang, Chenyang Wu, Tingting Wan, Shu Zhu

## Abstract

As the first-line barrier against food antigens, the intestinal epithelium’s role in establishing peripheral immune tolerance to food remains completely unknown. Additionally, the specific types of regulatory T cells induced to maintain food tolerance—along with their antigen specificity—are still unclear. Here we found small intestinal FOXP3^-^IL-10^+^ Tr1 cells, but not Treg cells, can respond to food antigens presented by intestinal epithelial cells (IECs) in an antigen-specific manner. Through immunopeptidome analysis of MHCII-presented peptides, we demonstrate that IECs can present food-derived antigens. Importantly, these IEC-presented food antigens are exclusively recognized by the TCR repertoire of small intestinal Tr1 cells, not Treg cells. Furthermore, we show that T cells recognizing IEC-presented food antigens specifically differentiate into Tr1 cells in the siLP. These findings establish two key insights: first, that maintaining food tolerance represents a physiological function of Tr1 cells, and second, that IECs play a central role in maintaining food tolerance. Our work thus provides mechanistic understanding of how the intestinal epithelium contributes to immune tolerance through its antigen presentation capacity and subsequent Tr1 cell induction.

## Introduction

The total surface area of the human small intestinal mucosa can reach 30 square meters and is continuously exposed to food-derived antigens. Adequate immune tolerance to food is required to prevent the unneccesary inflammatory responses to food antigens which is also consider “non-self” metarials^1, 2^. Intestinal professional antigen-presenting cells (pAPCs), including dendritic cells (DCs) and macrophages, have been shown to play pivotal roles in the establishment of immune tolerance to food antigens. CX3CR1^+^ macrophages resident in the lamina propria sampling food antigens in the intestinal lumen via transepithelial dendrites and transfer these antigens to CD103^+^ DCs^3^. CD103^+^ DCs migrate from the mucosa to the mesenteric lymph nodes (mLNs) and induce food antigen-specific FOXP3^+^ Tregs^4, 5^. Besides pAPCs, small intestinal epithelial cells also express MHCII and have been reported to shape the CD4^+^ T cell populations in the small intestine^6–9^. Our previous work has shown that lack of antigen presentation in IECs leads to food allergy^10^, but there is still no direct evidence that IECs can present food antigens and induce food antigen-specific T cells to mediate food tolerance in the small intestine.

CD4^+^ T cells with anti-inflammatory properties that originate from the mLN have been reported to mediate tolerance to food antigens^5, 11, 12^. It’s believed that FOXP3^+^ Tregs induced by CD103^+^ DCs can home to the siLP and mediate immune tolerance to food antigens^13, 14^. However, a recent study showed that most of the gliadin (a wheat protein)-responsive CD4^+^ T cells in the siLP are Th^lin-^ cells including FOXP3^-^ cells^11^. In addition to FOXP3^+^ Treg cells, FOXP3^-^IL10^+^ Tr1 cells constitute another regulatory T cell population in the small intestine. These cells populate the small intestine rather than the colon, consistent with the distribution of food antigens^15^. And the number of these FOXP3^-^IL10^+^ Tr1 cells shows periodic oscillations associated with daily food intake^7^. Although these evidence indicates potential correlation between Tr1 cells and food, and several pathological functions of Tr1 cells have been reported in colitis^16^, tumor^17^, T1D^18^ and EAE^19^ animal models, there is still lack of direct evidence that supports the physilogical function of Tr1 towards specific antigen such as food antigens.

Here, we demonstrate that small intestinal Tr1 cells, but not Treg cells, can respond to certain food antigens. By using immunopeptidomics to analyze peptides presented by IECs via MHCII, we found that IECs can present food protein derived antigens. These food antigens presented by IECs can only be recognized by the TCR of small intestinal Tr1 cells, but not Treg cells. These T cells that recognize food antigens presented by IECs specifically develop into Tr1 cells in siLP. Our work indentified maintaining food tolerance is a physilogical function of Tr1 cells, and revealed the key role of IECs in the establishment of food tolerance.

## Results

### Small intestinal Tr1 cells response to dietary antigen

In addition to FOXP3^+^ Treg cells, FOXP3^-^IL-10^+^ Tr1 cells constitute another CD4^+^ T cell subset with regulatory properties in siLP. Unlike Treg cells, which are distributed in multiple organs and tissues (lymphoid tissues and mucosal surface) (Extended Data Fig.1a-d), Tr1 cells specifically accumulate in the lamina propria of the small intestine among the 35 tissues and organs we examined (Extended Data Fig.1a-c, and e). Since food antigens are abundantly distributed in small intestine, and Tr1 cells decreased in GSDMD mutant mice in which food antigen can not induce MHCII expression in IECs^10^ as well as exhibit daily circadian rhytham due to the regular food intake^7^, we reasoned that while both Treg cells and Tr1 cells in small intestine may contribute to immune tolerance to food, and Tr1 cells may mainly reactive to food antigens.

We first evaluated the influence of the food antigens to these regulatory cell populations in IL-10^GFP^/FOXP3^RFP^ dual reporter mice. To our surprise, both the number and proportion of Treg cells in siLP did not change at all in mice fed an amino acid diet (AAD) specifically formulated to lack proteinaceous food antigens contraining majorly beta-caesin and kappa-caesin, however, Tr1 cells were found completely loss in the siLP of AAD-fed mice. Both Treg cells and Tr1 cells remained the same in the colon laminal propair (cLP) of AAD-fed mice (Fig. 1a,b and Extended Data Fig. 2a and b). These results led us to hypothesize that Tr1 cells rather than Treg cells respond to food-derived antigens in the small intestinal mucosa.

**Figure 1.**
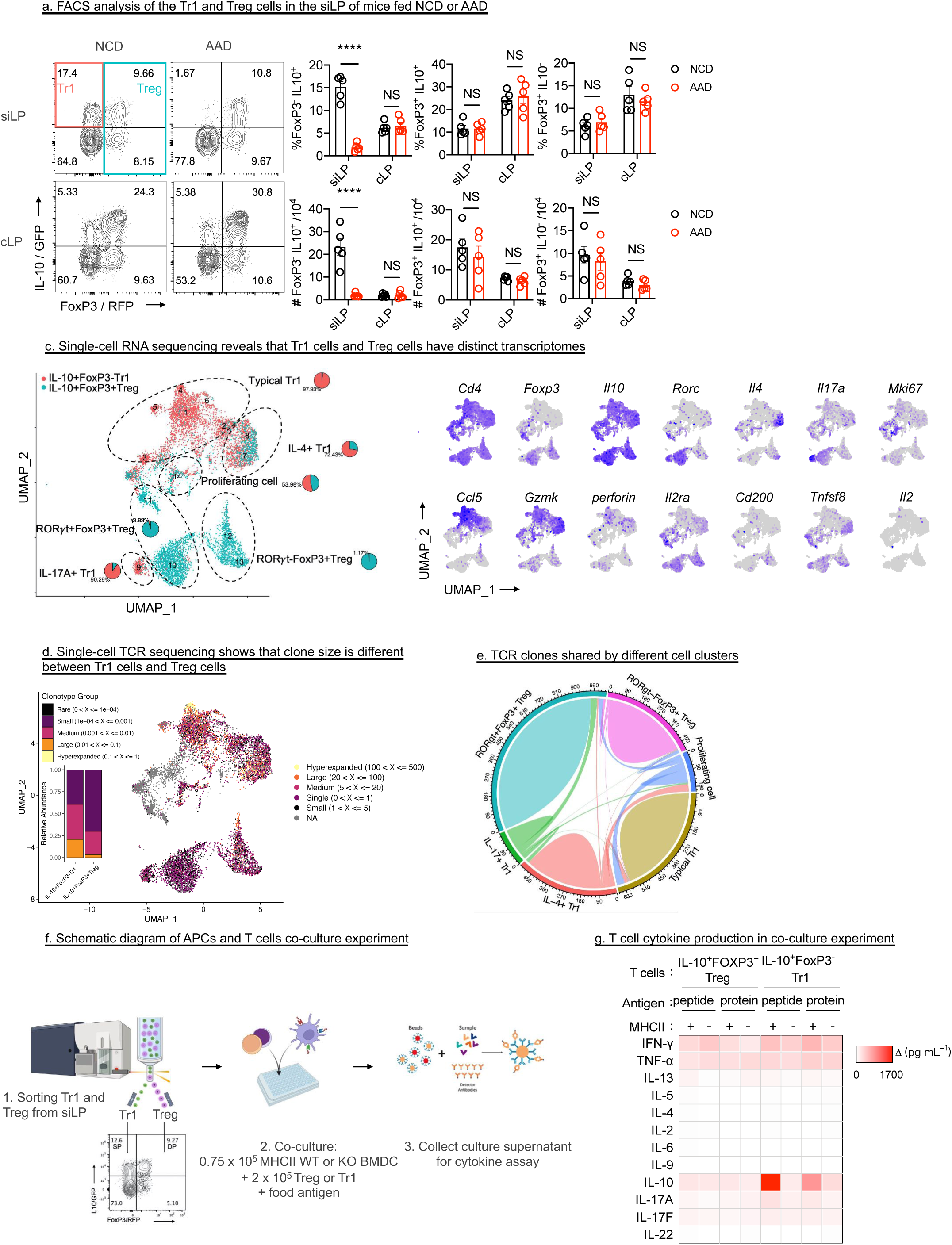
Tr1 but not Treg cells respond to dietary antigen in siLP. a, Representative FACS plots and quantification of FOXP3 and IL-10 expression on gated CD45^+^ CD3^+^ CD4^+^ T cells isolated from the siLP of mice fed the indicated diets (n=5). c-e, Single cell RNA and TCR sequencing of hashtag-labeled FOXP3^-^IL10^+^ Tr1 cells and FOXP3^+^IL-10^+^ Treg cells isolated from the siLP. (c) Uniform Manifold Approximation and Projection (UMAP) plot of the cells, with clusters annotated into 6 cell types (left). Single cell expression of the indicated genes projected on the UMAP plot (right). (d) TCR clonotype size distribution projected onto the UMAP plot. TCR clonotype abundance of Tr1 cells or Treg cells was quantified on a bar graph. (e) Circos plots showing TCR clonotypes shared between each annotated cell type. Each link shows how many T cells in that cell type have a shared TCR. f-g, FOXP3^-^IL10^+^ Tr1 cells or FOXP3^+^IL-10^+^ Treg cells were isolated from the siLP of NCD-fed mice and co-cultured with *H2-Ab1^+/+^*or *H2-Ab1^-/-^* bone marrow-derived DCs (BMDCs) in the presence of digested or intact casein proteins. (f) Schematic diagram of the co-culture experiment. (g) Measurement of T cell cytokine production in the co-culture experiment. Data are representative of at least three independent experiments (a, g). Data are mean ± s.e.m and were analyzed by two-way ANOVA Šídák’s multiple comparisons test, NS, no significance; \*\*\*\**P* < 0.0001.

We next sorted out Tr1 and Treg cells from the siLP of normal chow diet (NCD)-fed mice, label these two cell populations with hash tags and performed single cell RNA and TCR sequencing to compare the TCR repertoires between two cell populations. After demultiplexing, a total of 9,028 cells were recovered, of which 4,870 (53.94%) were Tr1 cells and 4,158 (46.06%) were Treg cells. The ratio of the two populations of cells is similar to that *in vivo* (Extended Data Fig. 1b). The cells were annotated based on their specific gene expression profiles and then clustered to 14 cell populations in an unsupervised manner (Fig.1c,d). 70.03% of the sorted Tr1 cells expressed *Il10*, *Ifng* and the marker of CD4+ T cell anergy^20^ as well as co-inhibitory molecules such as *Pdcd1* and *Lag3*, thus these cells are referred to as typical Tr1 cells (Fig.1c and Extended Data Fig. 3a). The remaining Tr1 cells co-expressed *Il10* and *Il17a* or co-expressed *Il10* and *Il4*, indicating that they may be in an intermediate state of transdifferentiation^21, 22^. Treg cells clustered into two groups based on the expression of *Rorc* (Fig.1c). Except for the proliferating cell cluster composed of both Tr1 cells (53.98%) and Treg cells (46.02%), the other cell clusters were either predominantly Tr1 cells (clusters 1-9) or predominantly Treg cells (clusters 10-13) (Fig.1c), indicating that these two cells populations exhibit obvious transcriptional differences. Compared with Treg cell subsets, typical Tr1 cell subsets have higher expression levels of *Ccl5*, *Gzmk* and *Prf1* (encode perforin) and lower expression levels of *Il2ra*, *Cd200* and *Tnfsf8*, which is consistent with the characteristics of Tr1 cells described in previous studies^17^.

We then analyzed the TCR repertoires in these two populations. One of the interesting finding is that the TCR clone size in typical Tr1 cells is much larger than that in Treg cells (Fig.1d and Extended Data Fig. 3b), indicating that the antigen pool recognized by typical Tr1 cells is more focused, whereas the antigen pool recognized by Treg cells is more diverse. Another surprising finding is that almost no TCR clones are shared by typical Tr1 cells and Treg cells (TCR clone sharing exists only in the proliferating cell population) (Fig.1e), and cells with the same TCR are aggregated in a specific cluster (Extended Data Fig. 4). These results suggested that the antigen specificity of Tr1 cells and Treg cells in siLP is completely different.

Since Tr1 cells and Treg cells in small intestine exhibit completely distinct TCR repertoire profiles, we next evaluated whether these two cell populations recoginze abundent antigen in the lumen of small intestine such as food antigens. We sorted Tr1 cells and IL-10^+^ Treg cells from the siLP of NCD-fed mice and co-cultured them with APCs loaded with dietary antigens that derived from casein protein, the main protein source in the mouse chow diet (Fig.1f and Extended Data Fig. 2a and b). In line with our previous finding of the loss of Tr1 cells in AAD-fed mice, we found Tr1 cells from NCD-fed mice, in contrast to Treg cells, can be stimulated by either digested or intact casein proteins (Fig.1g). In summary, these results demonstrated that Tr1 cells and Treg cells in siLP have distinct antigen specificities, and Tr1 cells is the main population that responding to food antigens.

### Tr1 cells recognize dietary antigen presented by IECs

According to our previous finding that MHCII induction in IECs is critical for immune tolerance to food^10^, we reasoned that food peptides-loaded MHCII in IECs might invovled in the generation of dietary antigen-specific Tr1 cells. We crossed *H2-Ab1*^Δ*IEC*^ mice with IL-10^GFP^/FOXP3^RFP^ dual reporter mice. The mice lacking MHCII in IECs resulted in an almost complete disappearance of Tr1 cells in the siLP under NCD-fed conditions (Fig.2a and b), phenotypically consistent with WT mice fed with AAD (Fig.1a), suggesting IEC antigen presentation is required for Tr1 cell development.

We next reasoned whether IECs can present food antigens to Tr1 cells. We performed mass spectrometry (MS) to evaluate the peptides bound to MHCII molecules that expressed on EpCAM^+^CD45^-^ IECs sorted from the small intestine of NCD-fed *H2-Ab1^f/f^* mice (Fig.2c). IECs sorted from NCD-fed *H2-Ab1*^Δ*IEC*^ mice or AAD-fed *H2-Ab1^f/f^* mice were used as negative controls (Fig.2c). The lengths of peptides identified in IEC MHCII immunopeptidome were normally distributed from 12∼20, and with a median length of 15-16 amino acids (Fig.2d and Extended Data Fig. 5a). More MHCII-bound peptides were found in IECs derived from NCD-fed *H2-Ab1^f/f^* mice than AAD-fed *H2-Ab1^f/f^* mice, MHCII-bound peptides were found in IECs derived from NCD-fed *H2-Ab1^f/f^* mice and that in NCD-fed *H2-Ab1*^Δ*IEC*^ mice were consider as the non-specific binding peptides, suggesting IEC MHCII present significant amount of antigens (Fig. 2d).

**Figure 2.**
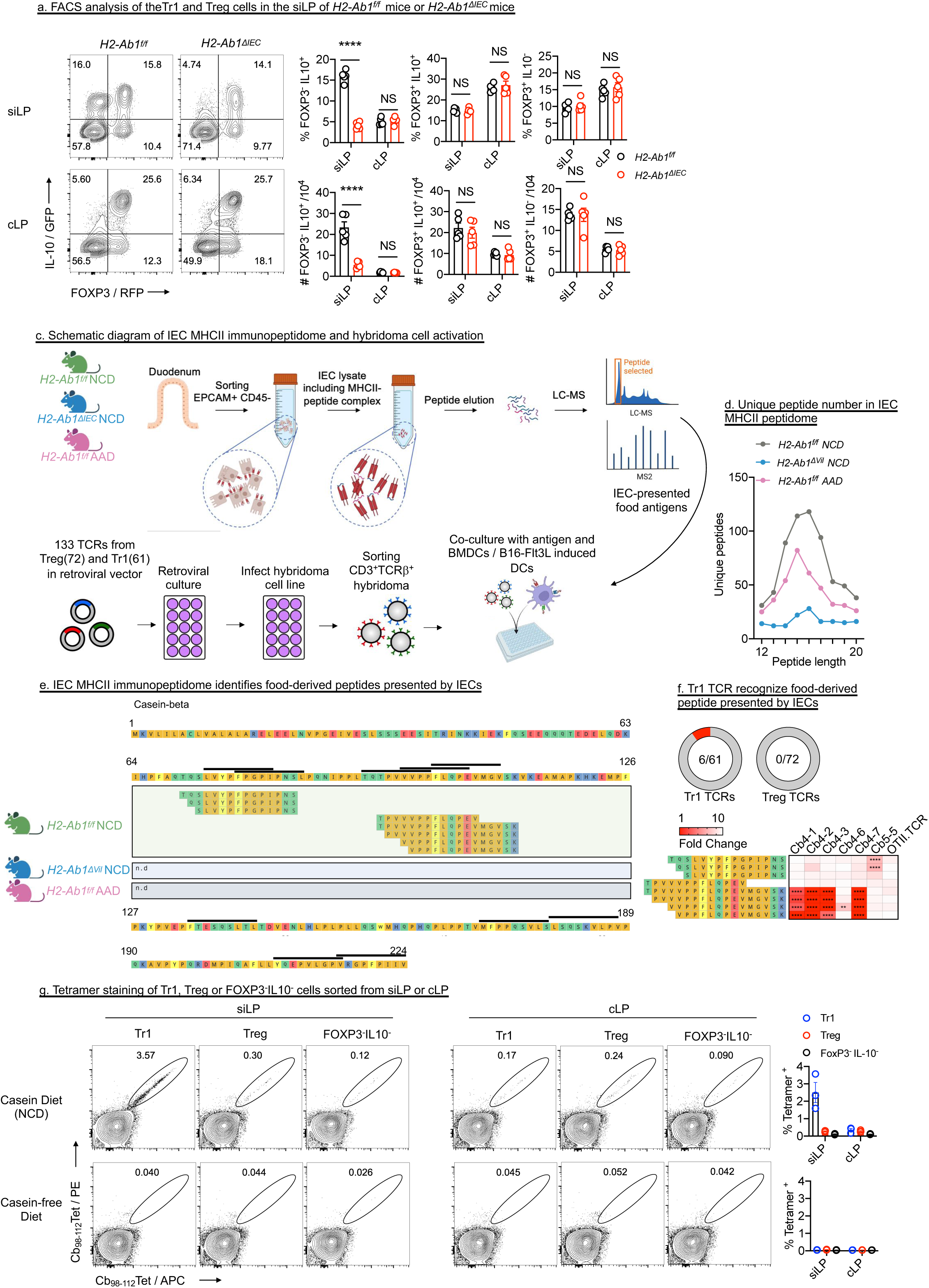
Tr1 cells recognize dietary antigen presented by IECs. a, Representative FACS plots and quantification of FOXP3 and IL-10 expression on gated CD45^+^ CD3^+^ CD4^+^ T cells isolated from the siLP of the indicated mice (n=5). c-f, The MHCII peptidome was used to identify antigens presented by IECs and screen the TCR repertoires of Tr1 cells and Treg cells for TCRs that recognize these antigens presented by IECs. (c) Schematic diagram of the experiment. MHCII peptidome was performed on IECs isolated from the indicated mice. TCRs from the TCR repertoires of Tr1 cells and Treg cells were cloned and expressed in 58α^-^β^-^ T cell hybridoma cell line. The hybridomas were then co-cultured with DCs loaded with the peptides identified in the IEC MHCII peptidome. (d) Length distribution of unique peptides identified in each MHCII peptidome sample. (e) Food protein-derived peptides identified in each IEC MHCII peptidome sample. The black lines above the protein sequences indicate predicted MHCII binding sites. (f) The number of TCRs in the TCR repertoires of Tr1 cells and Treg cells that recognize food antigens presented by IECs (upper panel). The response of 6 hybridoma cell lines carrying TCRs derived from Tr1 cells to each peptide (lower panel). g, Representative FACS plots and quantification of tetramer-positive Tr1 cells, Treg cells and FOXP3^-^IL-10^-^ cells sorted from the small or large intestine. Data were pooled from there independent experiments. Data are representative of three independent experiments (a). Data are mean ± s.e.m and were analyzed by two-way ANOVA Šídák’s multiple comparisons test, NS, no significance; \*\*\*\**P* < 0.0001.

The core binding motif of the MHCII-bound peptides from IECs derived from NCD-fed *H2-Ab1^f/f^*mice and AAD-fed *H2-Ab1^f/f^*mice, but not from NCD-fed *H2-Ab1*^Δ*IEC*^ mice, was consistent to previously indentified peptide motif that loaded by H2-Ab1 allele of typical APCs^23^ (Extended Data Fig. 5b), indicating that IEC-MHCII present similar peptides motif with typical APCs. Notably, 8 food-derived MHCII-bound peptides were detected only in the NCD-fed *H2-Ab1^f/f^* mice (Fig. 2e and Extended Data Fig. 5c). These peptides identified in the immunopeptidome are derived from two regions of the dietary casein, called Cb_70–84_ and Cb_95–112_ (Fig. 2e), which are also predicted to have strong binding affinities for MHCII molecules^23, 24^. Thus, we identified the casein peptides that can be presented by IEC-MHCII.

We next investigated whether food antigens presented by IECs could be recognized by the TCR characterized by the aforementioned single cell TCR sequencing data set. We retrieved paired TCR sequences from 133 most expanded Tr1 cell or Treg cell clones (Fig. 1c). We then cloned these TCRs into NFAT-GFP 58α^-^β^-^ T cell hybridoma cell line and tested their response toward 8 peptides identified in the aforementioned IEC MHCII immunopeptidome (Fig. 2c-e). Among the 61 TCRs from typical Tr1 cells, 6 TCRs could respond to the peptides mixture; however, none of the 72 Treg TCRs responds to the peptides mixture (Fig. 2f and Extended Data Fig. 4a). We further tested these 8 epitopes recognized by these 6 Tr1 TCRs one by one. 1 TCRs had relatively weak responses to the Cb_70–84_ region, and the rest of 5 TCRs responded to the Cb_95–112_ region, with the Cb4-7 TCR having the strongest response to Cb_98-112_ (Fig. 2f and Extended Data Fig. 4b).

Thus, we generated MHCII tetramers loaded with the Cb_98-112_ peptide (Cb_98-112_ tetramer) to stain casein-specific CD4^+^ T cells *in vivo*. We sorted equal numbers of polyclonal Tr1, Treg and FOXP3^-^IL10^-^ cells from the siLP of NCD-fed IL-10^GFP^/FOXP3^RFP^ dual reporter mice and stained these cells by the Cb_98-112_ tetramer (Extended Data Fig. 6a, b). The Cb_98-112_ tetramer stained significant amount of Tr1 cells isolated from siLP but almost no Treg cells or FOXP3^-^IL10^-^ cells (Fig. 2g), which is consistent with the previous findings that it is small intesinal Tr1 cells rather than Treg cells that respond to dietary casein. Since feeding mice with a diet lacking proteinous antigens results in loss of MHCII expression by IECs and disappearance of Tr1 cells^10^, we used a casein-free diet (CFD) as a control, which contains intact food antigens derived from whey and the MHCII expression on IECs and Tr1 cells in the siLP are at the same levels with the mice fed with the casein diet (NCD) (Extended Data Fig. 6c-e). Tr1 cells in the siLP from mice fed a CFD were completely unstained with the Cb_98-112_ tetramer (Fig. 2g), confirming the existence of casein-specific Tr1 cells in the small intestine. Taken together, these results demonstrated that Tr1 cells rather than Treg cells respond to certain food antigens presented by IECs in siLPs.

### Food-specific T cells developed into Tr1 cells in siLP

Given that Tr1 cells specificly recognize certain food antigens such as casein, we then investigate how food antigen-specific naïve T cells develop into Tr1 cells in siLP. To this end, we generated the transgenic mice carring Cb4-7 TCR (Cb4-7Tg), which give strongest response toward casein Cb_98-112_ antigen presented by IECs. Then Cb4-7Tg were crossed with IL-10^GFP^/FOXP3^RFP^ dual reporter mice to analyze the antigen-specific Tr1 cells. Cb4-7Tg T cells can be specifically stained by the Cb_98-112_ tetramer but not OVA specific tetramer, and the staining efficiency of Tr1, Treg and FOXP3^-^IL10^-^ cells isolated from the siLP of Cb4-7Tg mice does not differ significantly (Extended Data Fig. 7a), confirming that the transgenic Cb4-7 TCR is uniformly expressed in different T cell sub-populations. Naïve T cells isolated from Cb4-7Tg mice were transferred into recipient mice fed with NCD that containing intact casein protein or AAD without peptides digested from casein (Extended Data Fig. 7b). Donor Cb4-7Tg T cells accumulated more in the siLP of NCD-fed mice (Extended Data Fig. 7c). We found that half of the donor T cells developed into Tr1 cells in the siLP of mice fed NCD but not AAD (Extended Data Fig. 7d-e). Since IECs from AAD-fed mice lacks both casein antigen and MHCII expression (cite cell paper), it’s still not clear whether the generation of Tr1 cells in the siLP requires the cognate food antigen.

Thus we isolated naïve T cells from Cb4-7Tg mice and then transferred them into recipient mice fed with NCD (with casein antigen) or CFD (with whey antigen but without casein antigen) (Fig. 3a). IECs from mice fed with both diets expressed MHCII and have Tr1 cells in siLP (Extended Data Fig. 6c-e). Donor Cb4-7Tg T cells acquired the Tr1 phenotype in the siLP, but not in the cLP, of the mice fed with NCD (Fig.3b,c). Although the receipint mice fed with CFD or NCD showed the same number of Tr1 cells in steady state, the donor Cb4-7Tg T cells largely become Tr1 cells in receipint mice fed with NCD but no Cb4-7Tg T cells become Tr1 cells at all in receipint mice fed with CFD (Fig.3b, c and Extended Data Fig. 7f). These results indicate that the generation of food antigen-specific Tr1 cells in the siLP requires the presence of the cognate food antigen.

**Figure 3.**
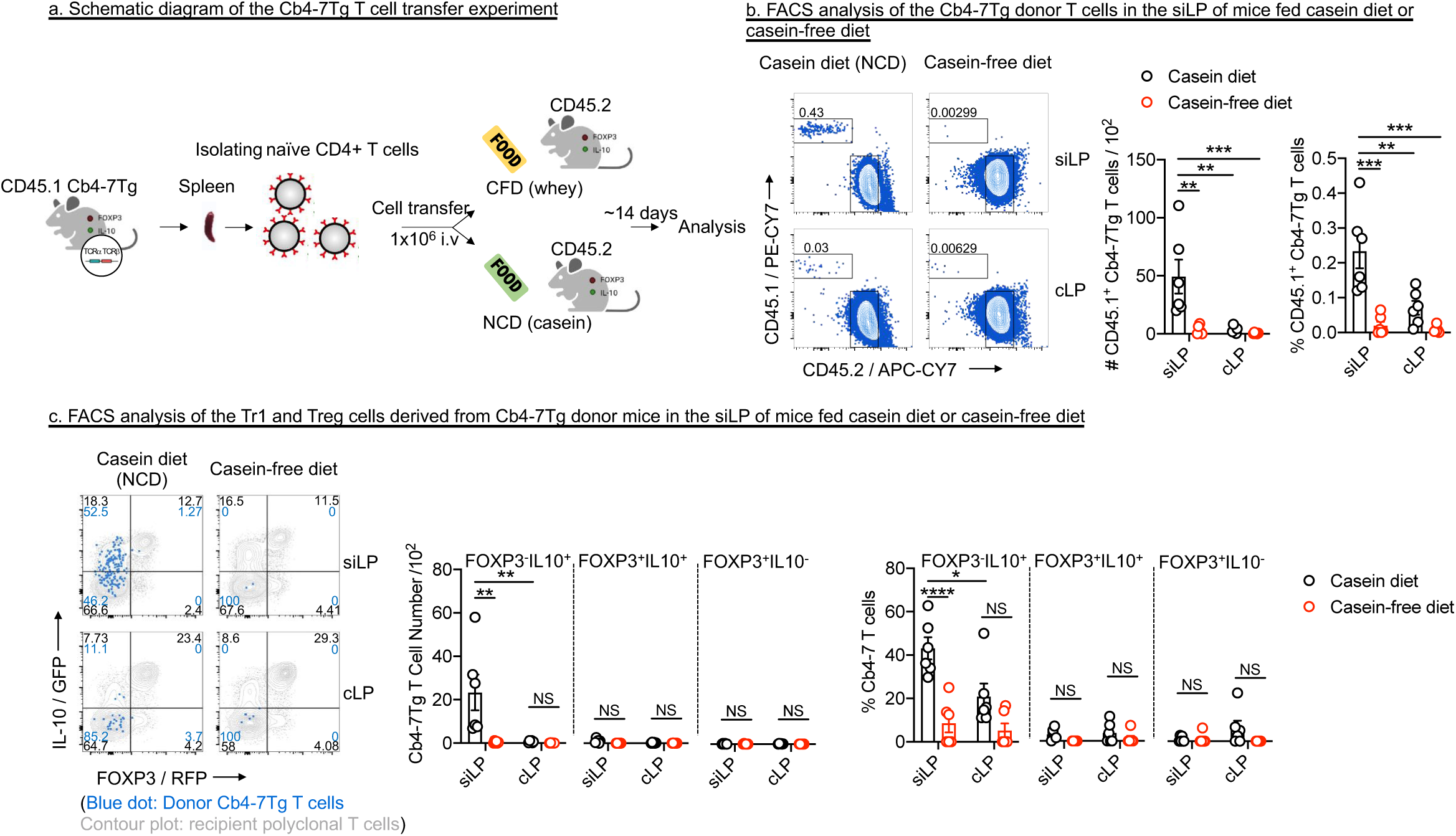
Food-specific T cells developed into Tr1 cells in siLP. a-c, CD45.1^+^ Cb4-7 Tg T cells were transferred into CD45.2^+^ hosts fed NCD or CFD (n=5). (a) Experimental schematic. (b) Representative FACS plots and quantification of donor CD45.1^+^ Cb4-7 Tg T cells in siLP or cLP of host mice. (c) Representative FACS plots and quantification of FOXP3 and IL-10 expression in donor (CD45.1^+^) and host polyclonal CD4^+^ T cells from siLP or cLP. Data are representative of three independent experiments. Data are mean ± s.e.m and were analyzed by two-way ANOVA Šídák’s multiple comparisons test, NS, no significance; \**P* < 0.05, \*\**P <* 0.01, \*\*\**P* < 0.001, \*\*\*\**P* < 0.0001.

## Discussion

Although many studies have reported the functions of Tr1 cells in pathological conditions of various immune-related diseases, their physiological responses to specific antigens remain unclear^22^. Here, we found that Tr1 cells in siLP can respond to food antigens presented by IECs and mediate immune tolerance to food antigens.

Both mice and humans have two regulatory T cell subsets, Tr1 and Treg, with completely different transcriptional characteristics. Unlike Treg cells, which have a defined developmental process, the generation of Tr1 cells is affected by a variety of external stimuli and can develop from memory T cells or transdifferentiate from effector Th1, Th2, and Th17 cells^21, 22^. Therefore, Tr1 cells appear to be a more plastic regulatory T cell subset. Our data also support this point: During in vitro culture, Treg cells isolated from the small intestine maintain stable IL-10 expression, while Tr1 cells gradually downregulate IL-10 unless cultured with antigen-presenting IECs.

The microenvironment-sensitive nature of Tr1 cells makes them more likely to mediate transient immune tolerance. Food antigens recognized by Tr1 cells only appear after eating, whereas the self-antigens or commensal microorganism antigens recognized by Treg cells persist continuously^25^. Thus, Tr1 and Treg cells may mediate immune responses to antigens with different occurrence frequencies.

IECs have been reported to constitutively express MHCII and shape intestinal CD4^+^ T cell population^6–10^, but the specific antigens presented by IECs through MHCII under physiological conditions had not been identified. Here, we performed mass spectrometry to analyze the epitopes bound to MHCII molecules on IECs. We found that food-derived casein peptides were presented by IECs and presentation of food antigens by IECs induces food antigen-specific Tr1 cells in the siLP in steady state. Notably, food antigen-specific T cells differentiate into Tr1 cells after migrating to the intestine and receiving food antigen presentation by IECs.

IECs cover the entire intestinal mucosa and consequently present vast amounts of antigen. Previous studies demonstrate that Tr1 cell generation depends on cognate antigen dose, with high concentrations of MHCII-restricted antigens promoting Tr1 cell development^17^. When food antigen-specific T cells migrate to the small intestine, they encounter abundant food antigens presented by IECs – this likely explains why IECs specifically induce Tr1 cell differentiation in the small intestine^10^.

## Acknowledgement

We thank Mo Xu from National Institute of Biological Sciences for providing the vectors used to produce the TCR transgenic mice, the 58α^-^β^-^ T cell hybridoma cell line, and the MIGR1-Thy1.1 plasmid. We thank Jincun Zhao from Guangzhou Medical University for the assistance provided during the production of TCR transgenic mice. We thank Rongbin Zhou for providing *Vil-Cre*, *CD11c-gfp-Cre* mice. We thank Wen Pan for providing *Vil-Cre-ER^T^*^2^ mice. We thank Mengjie Liu and Hong Yao for their help in animal experiments. We thank Jian Wang and members of his laboratory for advice and assistance with T cell factor assays. We thank Hai Li and members of his laboratory for assistance with TCR cloning. This work was supported by grants from the National Natural Science Foundation of China (82325025) (SZ), the National Key R&D Program of China (2023YFC2306101) (SZ), the CAS Project for Young Scientists in Basic Research (YSBR-074) (SZ).

## Author Contributions

S.Z., T.Z., and G.Z. conceived the project and designed the experiments. T.Z., G.Z. and C.W. performed the experiments and collected the data. T.Z. analyzed the scRNA-seq and bulk RNA-seq data. T.W.bred the IL-10^GFP^/FOXP3^RFP^ dual reporter mice. S.Z., T.Z., and G.Z. wrote the manuscript. S.Z. supervised the project.

## Methods

### Mice

IL-10^GFP^/FOXP3^RFP^ dual reporter mice (IL-10^GFP/+^ mice crossed with FOXP3^RFP^ mice) and CD45.1 mice were kindly provided by Prof. Richard A. Flavell of Yale university school of medicine. *Vil-Cre*, *CD11c-gfp-Cre* mice were provided by Dr. Rongbin Zhou of University of Science and Technology of China (USTC). *Cd19-Cre* mice were purchased from GemPharmatech (Nanjing, CN). Cb4-7 TCR transgenic mice was generated in our lab. *Vil-Cre-ER^T^*^2^ mice were provided by Dr. Wen Pan of USTC. Sex– and age-matched littermates aged of 8-10 weeks were used in most of the studies except for those in diet switch experiment (Sacrificed at 12 weeks old) or those fed AAD (weaning). All mice were kept under specific pathogen-free (SPF) conditions under a strict 12-h light cycle (lights on at 08:00 and off at 20:00) in the animal facility at USTC. All experimental procedures carried out in mice were with reference to the National Guidelines for Animal Usage in Research (China) and were approved by the Animal Ethics Committee of USTC (USTCACUC212101033).

### Diet

The AAD was customized by Trophic diet (Jiangsu, CN). The ingredients of AAD were listed as follows (g/kg): L-Alanine 4.5; L-Arginine, 6.3; L-Aspartic Acid 11.3; L-Cystine 3.7; L-Glutamic Acid 36.2; Glycine 3.1; L-Histidine, 4.5; L-Isoleucine 8.4; L-Leucine 15.3; L-Lysine-HCl 16.1; L-Methionine 4.5; L-Phenylalanine 8.7; L-Proline 20.4; L-Serine 9.4; L-Threonine 6.6; L-Tryptophan 2.1; L-Tyrosine 9.2; L-Valine 9.9; Sucrose 100; Cornstarch 399.886; Dyetrose 145; Soybean Oil 70; tBHQ 0.014; Cellulose 50; Salt Mix 35; Sodium Bicarbonate 7.4; Vitamin Mix 10; Choline Bitartrate 2.5. In the second week after birth, the diet in the breeding cages was replaced with AAD, and after weaning, the mice were fed AAD until they were sacrificed. Casein diet (also used as NCD in this paper) and casein-free diet are also purified diets that purchased from Trophic diet (Jiangsu, CN). The composition of these diets is identical to AAD, except that the amino acids are replaced by an equivalent amount of intact proteins. All diets were irradiated, vacuum-packed and stored at –20°C. The diets placed in the mouse cages were changed twice a week. All mice had *ad libitum* access to food.

### Antibodies, Flow cytometry and cell sorting

Single-cell suspensions were incubated on ice with conjugated antibodies in MACS buffer (PBS containing 2% FBS and 1 mM EDTA). Unlabelled anti-CD16/32 (clone 93, Biolegend) was used to block Fc receptors. Dead cells were excluded with Zombie Aqua™ dye (Biolegend) or DAPI (3 μM, Biolegend). The following antibodies were used for mouse cell-surface staining: CD4 PE/Cy7 (GK1.5) Biolegend 100422, CD45 PerCP/Cy5.5 (30-F11) Biolegend 103132, CD4 PerCP/Cy5.5 (GK1.5) Biolegend 100434, CD45.1 PE/Cy7 (A20) Biolegend 110730, CD45.2 PE/Cy7 (104) Biolegend 109830, CD3 APC/Cy7 (17A2) Biolegend 100222, CD4 APC (GK1.5) Biolegend 100412, CD49b APC (HMα2) Biolegend 103516, LAG-3 PE/Cy7 (C9B7W) Biolegend 125226, CD90.1 PE (OX-7) Biolegend 202524, TCRβ APC (H57-597) Biolegend 109212, CD3 PE/Cy7 (17A2) Biolegend 100220, CD4 Super Bright 600 (RM4-5) Invitrogen 63-0042-82, CD45.2 APC/Cy7 (104) Biolegend 109824, I-A/I-E FITC (M5/114.15.2) Biolegend 107606, CD326 (EpCAM) APC (G8.8) eBioscience 17-5791-82, CD11c PE (N418) Biolegend 117308, CD11c PerCP/Cy5.5 (N418) Biolegend 117328, CD19 APC/Cy7 (6D5) Biolegend 115530. Except for CD49b and LAG-3, which were stained at 37°C for 30 minutes, all other surface markers were stained at 4°C for 15 minutes. The following antibodies were used for mouse intracellular staining: Tbet PE/Dazzle™ 594 (4B10) Biolegend 100456, GATA3 PE (L50-823) BD 560068, RORγt BV650 (Q31-378) BD 564722, FOXP3 Alexa Fluor 647 (MF-14) Biolegend 126408, IFN-γ PE/Cy7 (XMG1.2) Biolegend 505826, IL-17A APC (TC11-18H10.1) Biolegend 506916, IL-4 PE (11B11) Biolegend 504104.

For intracellular transcription factor staining, cells were fixed/permeabilized for 30 minutes at room temperature or overnight at 4°C with FOXP3/Transcription Factor Staining Buffer Set (eBioscience). Then the cells were washed with 1x Permeabilization solution (eBioscience) and stained with fluorescently-conjugated antibodies in 1x permeabilization buffer for 30 minutes at room temperature. Stained cells were washed three times with 1x PBS. For intracellular cytokine staining, cells were first incubated for 4 hours in RPMI 1640 medium with 10% FBS, Cell Stimulation Cocktail (Invitrogen) at 37°C. Then cells were fixed/permeabilized for 20 minutes at 4°C with Fixation/Permeabilization Solution Kit (BD) and then washed with 1x Permeabilization solution (BD) and stained with fluorescently-conjugated antibodies in 1x permeabilization buffer for 30 minutes at 4°C. Stained cells were washed three times with 1x PBS. Flow cytometry data were collected using CytoFlex S (Beckman Coulter) or Cytek Aurora (Cytek) and analysed with FlowJo V10 software (Tree Star). Lymphocytes were sorted using FACSAria Fusion (BD) or CytoFlex SRT (Beckman Coulter). IECs were sorted by CytoFlex SRT using vertical sorting mode.

For analysis of cell numbers, the number of cells per microliter was obtained from the Cytoflex S. This number was first multiplied by 200 to obtain the total number of cells per tube and then multiplied by 5 to obtain the total number of cells in the intestinal segment, since only one fifth of the cells were stained for flow cytometry analysis. For the small intestine, the total number of cells in the whole small intestine was multiplied by 4 based on the total number of cells in the intestinal segment, since only one quarter of the tissue was taken for digestion due to the length of the small intestine (one quarter each of the duodenum, jejunum, and ileum).

### Cell isolation from the intestine

Mice were lethally anesthetized with pentobarbital sodium and perfused with PBS containing 2.5% FCS. For the isolation of IECs and intraepithelial lymphocytes, intestinal tissues were excised and flushed thoroughly with ice old PBS. Use ophthalmic forceps to remove the fat tissue on the surface of the intestine under a stereomicroscope. The intestines were then turned inside out and cut into 1 cm sections then transferred into RPMI with 1.5 mM EDTA (Sigma-Aldrich) and 2% FCS, and shaken for 15 min at 37 °C. Supernatants were collected through a 100-mm cell strainer to get single-cell suspensions containing both epithelial cells (∼90%) and lymphocytes (IEL, ∼10%).

For the isolation of lamina propria lymphocytes, intestinal tissues were excised and flushed thoroughly with ice old PBS. Use ophthalmic forceps to remove the fat tissue on the surface of the intestine under a stereomicroscope. The intestines were then turned inside out and cut into 1 cm sections then transferred into RPMI with 1.5 mM EDTA, 1mM dithiothreitol (Sigma-Aldrich) and 2% FCS, and shaken for 15 min at 37 °C for 3 times. Then the supernatants were discarded. Collect tissue sections and washed with RPMI with 3% FCS (EDTA-free) for another 5 min. The intestinal sections were transferred into digestion buffer (RPMI with 0.5 mg/ml collagenases II (Sigma-Aldrich), 0.5 mg/ml Dnase I (Roche) and 5% FBS (Gibco)) and shaken for 35 min at 37 °C. Stop the digestion by adding EDTA to the digestion buffer. Supernatants were collected through a 100-mm cell strainer to get single-cell suspensions and lymphocytes were enriched by 40% Percoll (Cytiva) gradient centrifugation.

### Tissue digestion

Mice were lethally anesthetized with pentobarbital sodium and perfused with PBS containing 2.5% FCS. Tissues were harvested into HBSS with 2.5% FCS and 2mM EDTA, and stored on ice until further processing. The thymus, lymph nodes, Peyer’s patches, spleen, adrenal glands were dispersed with frosted glass microscope slides, filtered through 100mm mesh. Where necessary, erythrocytes were lysed prior to counting using a Countess Automated Cell Counter (ThermoFisher). The liver was prepared similarly, except the resulting cell suspension was washed extensively and centrifuged (600g, 10min) through a 40% Percoll (Sigma Aldrich) solution prior to erythrocyte lysis. Bone marrow was extracted from a single femur by crushing with a mortar and pestle. Blood was treated to lyse red cells prior to staining.

The salivary glands, lungs, pancreas, kidney, adipose tissue, reproductive tissues, eyes, back skin, tongue, hind limb muscle, heart, bladder and brain were digested as follows to extract tissue leukocytes: Tissues were chopped finely with razor blades and washed by centrifugation to remove debris. They were then resuspended in digest buffer (IMDM supplemented with 20% FCS, 10mM HEPES, 1mM sodium pyruvate, 10mg/ml gentamicin, 1mM CaCl2 and 1mM MgCl2) with enzymes (400mg/ml collagenase IV, 100mg/ml hyaluronidase and 40mg/ml DNase I, Sigma Aldrich). In some experiments 2mg/ml collagenase IV was used in place of collagenase D. Tissues were shaken in an agitating mixer at 37° for 15min, dispersed with a pipette, and shaken for another 15min. The solution was then filtered through 100mm mesh and any remaining chunks were forced through the mesh. The cells were washed and passed through 40% Percoll to enrich for the leukocyte fraction.

### Single cell RNA and TCR sequencing

We used hashtag reagents for sample barcoding, enabling the amalgamation of two samples into a single lane for subsequent demultiplexing during analysis. Specifically, the hashtags consisted of two antibodies recognizing ubiquitous surface markers, CD45 and MHC class I, each conjugated to the same oligonucleotide containing the barcode sequence.

Lamina propria lymphocytes were isolated from the proximal small intestine of IL-10^GFP^/FOXP3^RFP^ dual reporter mice and Tr1 and Treg cells were sorted. Cell viability and integrity were evaluated by Trypan blue staining. Subsequently, cells were incubated with 1 µl (0.5 µg) of the respective Totalseq-C (Hashtag1 for Tr1 cells and Hashtag2 for Treg

cells) anti-mouse Hashtag antibodies (155861 and 155863, BioLegend) for 30 min at 4 °C. After staining, the samples were washed twice with 500µl of cell staining buffer (BioLegend) and pooled into a single tube. The ratio of Tr1 and Treg cells was not adjusted. Finally, the cells were re-suspended in a resuspension buffer (PBS containing 0.04% BSA). The cell number and viability were evaluated again, and the optimal cell concentration (700 – 1200 cells/µl) was set according to the 10x Genomics protocol.

Cells were loaded onto the 10X Chromium Single Cell Platform (10X Genomics) at a concentration of 1,000 cells per µl (Single Cell 5’ library and Gel Bead Kit v.2) as described in the manufacturer’s protocol. Generation of gel beads in emulsion (GEMs), barcoding, GEM-RT clean-up, complementary DNA amplification, gene and TCR, hashtag library construction were all performed as per the manufacturer’s protocol. Qubit was used for library quantification before pooling. The final library pool was sequenced on the Illumina Novaseq 6000 instrument using 150-base-pair paired-end reads.

Gene expression count matrices were generated using Cell Ranger version 5.0 count method using the default parameters. Sequencing files were aligned to the refdata-gex-mm10-2020-A reference library provided by 10X Genomics. Raw outputs from Cell Ranger were processed in the Seurat R package (4.1.1). hashing-based doublet detection strategy HTODemux was used to identify doublets that represent two or more cells representing different samples (positive.quantile = 0.99). Genes whose sum of expression in all cells was less than 3 were deleted. We retained cells that had unique feature counts within range from 200 to 5000 and mitochondria counts of less than 10%, log-normalized the counts and scaled each gene. 2000 highly variable features were determined using the “vst” method (FindVariableFeatures function), and served as input to the principal component analysis for dimensionality reduction. We used the first 25 principal components to cluster cells using the Louvain algorithm. To determine cluster specific transcriptional programs, differential gene expression analysis was performed with Wilcoxon Rank Sum Test.

Single cell TCR sequencing data was assembled and the clonotypes were determined using the default settings of Cell Ranger version 5.0 VDJ pipeline. combineTCR and combineExpression function in scRepertoire (2.0.0) package was used to merge scTCR-seq data with the Seurat object of scRNA-seq data. the relative number of clonotypes shared between each cell type was calculated using the getCirclize function. The Circos plot was generated by R package circlize (0.4.16).

### IEC MHCII peptidome

Epithelial cells from the proximal small intestine of the same mice that underwent single-cell RNA and TCR sequencing were used for MHCII-associated peptide isolation. IECs were sorted by CytoFlex SRT using vertical sorting mode. Approximately 10^7^ cells were lysed in immunoprecipitation buffer (YUBiomics). MHCII-antigen complexes were captured using Protein G Magnetic Beads (Promega) pre-coated with anti-mouse MHC class II (I-A/I-E) antibody (BioXcell) according to the YUBiomics NeoAntigen Platform protocol. Following immunoprecipitation, bound complexes were eluted and peptides were dissociated by acid treatment. The eluted peptides were filtered through 10-kDa molecular weight cutoff filters and lyophilized prior to mass spectrometry analysis.

Mass spectrometric analysis was performed using a timsTOF Pro mass spectrometer (Bruker Daltonics) coupled to an Evosep One liquid chromatography system (Evosep). Peptides were separated using a 21-minute gradient method. The mass spectrometer was operated in data-dependent acquisition mode with the following parameters: MS1 scan range of 100-1700 m/z in positive ion mode; accumulation and ramp times of 100 ms each; dynamic exclusion duration of 24 seconds; ion source voltage set to 1500 V; source temperature maintained at 180°C; and dry gas flow rate of 3 L/min. Ion mobility separation was conducted with a mobility range of 0.75-1.35 Vs/cm^2^, followed by acquisition of 8 PASEF MS/MS scans per cycle.

The acquired raw data were processed using PEAKS X Pro software (version 10.6, Bioinformatics Solutions Inc.) with searches against both the UniProt mouse reference proteome database, food-derived protein database and proteome database. Peptide identification was performed with a false discovery rate threshold of <1% at the PSM level.

### Co-culture of DCs and T cells and Cytokine measurements

2×10^5^ sorted T cells were co-cultured with 1×10^5^ BMDCs in the presence of intact casein protein (Sigma, C8654) or digested casein protein (50ug/mL). The in vitro protein digestion method was modified from a previously reported method. In brief, after the protein suspension temperature was stabilized at 37°C, the pH was adjusted to 8.0 using NaOH or HCl. Trypsin (Sigma, T0303) at a working concentration of 0.1 mg/mL was added to the protein suspension and incubated for 5 minutes. Then an equal amount of trypsin inhibitor (Sigma, T9128) was added. Cell culture supernatants were harvested after 24 h of co-culture and stored at –20°C until the cytokine assays. The concentration of IFN-γ,TNF-α, IL-13, IL-5, IL-4, IL-2, IL-6, IL-9, IL-10, IL-17A, IL-17F and IL-22 in supernatants was measured by The Mouse Th Cytokine Panel (12-plex) LEGENDplex Multi-analyte Flow Assay kit (BioLegend, 741044) according to the manufacturing protocol. CytoFlex S (Beckman Coulter) was used to analysis the beads in the cytokine kit. The fcs data were analyzed by an online software (https://www.biolegend.com/enus/immunoassays/legendplex/support/software) provided by manufacturer.

### BMDCs

BMDCs from C57BL/6 mice were cultured on 10 cm non-treated cell culture Petri dishes in RPMI with 10% FBS, 20 ng/mL GM-CSF (Biolegend, 576308) and 100 U/mL penicillin– streptomycin (Gibco, 10378016). Every 1–2 days for the first 4 days, plates were gently washed and non-adherent granulocytes removed by aspirating 50% of the culture media with subsequent replacement of fresh media. On day 4, media was aspirated completely and replaced with fresh culture media with 20 ng/mL GM-CSF. On day 6, BMDC plates were washed with PBS and loosely adherent and non-adherent cells collected. Cells were centrifuged at 500g for 5 min, resuspended in fresh culture media and replated on 10 cm non-treated cell culture Petri dishes. On days 7–8, plates were washed with PBS and loosely adherent and non-adherent cells were collected.

### TCR cloning

Partial sequences of the most abundant TCRs in the TCR repertoires of Tr1 cells and Treg cells was obtained from single-cell TCR sequencing data. Tcrdist3 (0.1.0) was used to calculate distance between each TCR to remove highly similar TCRs in our TCR sequencing data. Full-length sequence of the TCR was obtained using the V-QUEST tool and Gene database provided by IMGT (https://www.imgt.org/).

To reconstitute TCRs, cDNA of TCRα and TCRβ were linked with the self-cleavage sequence of 2A (TCRα-p2A-TCRβ), and shuttled into a modified MIGR1 retrovector in which IRES-GFP was replaced with IRES-Thy1.1. Briefly, The target DNA sequence and the DNA oligo primers were synthesized from Qingke (For Cb4-7 TCR: forward,5’-GGCGCCGGAATTAGATCTCTCGAGGCCACCATGGACAAGATCCTGA-3’; reverse, 5’-AACGTTAGGGGGGGGGGGCGGAATTCAGGAATTTTTTTTCTTGACCAT-3’). The target DNA fragment was amplified by PCR using 2× Phanta Max Master Mix (Vazyme, P515-02). The vector plasmid was digested with XhoⅠ (NEB, R0146S) and EcoRⅠ (NEB, R3101S). The DNA fragments were size-verified by gel electrophoresis. The DNA fragments were purified with Gel Extraction Kit (CWBIO, CW2302M) following the manufacturer’s instructions. The DNA fragments were assembled using Exnase MultiS (Vazyme, C113-01). 5 µL of the recombinant plasmid was added to 50 µL of DH5α Competent E. coli Cells (Qingke, DLC101) and incubated on ice for 30 min. The mixture was heat-shocked at 42°C for 60 s, followed by incubation on ice for 5 min. The bacterial suspension was spread onto Amp+ LB plates and incubated at 37°C for 12 hrs. single colony was picked and subjected to PCR amplification for verification using Rapid Taq Master Mix (Vazyme, P222-01). The correct single colony was inoculated into 15 mL of liquid LB medium and cultured overnight in a 37°C shaking incubator. The bacterial culture was centrifuged at 3000 rpm for 10 min and plasmid DNA was extracted using the Plasmid Midi Preparation Kit (TIANGEN, DP106-02) the following day.

### Generation of TCR hybridomas

8 μg of TCR-MIGR1-Thy1.1 plasmid and 8 μg of pCL-Eco plasmid were added to Opti-MEM (total volume of 500μL) in a sterile tube and gently pipetted. 64 μg PEI (2 μg/mL company) was added to 468 μL Opti-MEM and mixed thoroughly. The PEI/Opti-MEM mixture was added dropwise to the DNA/Opti-MEM mixture, mixed by pipetting, and incubated at room temperature for 15 min. The PEI/DNA mixture was added dropwise to 293T cells at 50–70% confluency. The plate was gently swirled and incubated at 37°C for 5 hr. The medium was replaced with fresh complete DMEM and incubated at 37°C for 24 hr. The DMEM was replaced with T cell culture medium (TCM). TCM was prepared by supplementing 445 mL RPMI-1640 (Gibco, 11875093) with 50 mL FBS, 5 mL penicillin-streptomycin (Gibco, 10378016), and 500 μL of 55 mM 2-mercaptoethanol (Gibco, 21985023). 293T were cultured for an additional 24 hr. TCM containing virus was then collected.

Polybrene was added to the collected TCM (8 μg/mL). NFAT-GFP 58α-β-hybridoma cells (1×10⁶) were resuspended in 2 mL of TCM containing 8μg/mL polybrene. The cells were plated in a 6-well plate and centrifuged at 2000g for 2 hr at 33°C. The cells were gently resuspended by pipetting and incubated at 37°C for 2 hr. Cells were centrifuged at 800 rpm for 5 min at 4°C and cultured in 2 ml of fresh pre-warmed TCM medium for 48 hr.

Then the hybridomas were transferred to a 15 ml Falcon tube and centrifuged at 800 rpm for 5 min at 4°C. The pellet was resuspended in 200μL antibody mixture (0.8μL P7-CD4 (GK1.5, Biolegend, 100422), 0.8μL APC-TCRβ (H57-597, Biolegend, 109212) in 200μL

MACS buffer) and incubated for 15 min at 4°C. Hybridomas were washed with 5 mL PBS and centrifuged at 800 rpm for 5 min at 4°C. The pellet was resuspended in 500μL MACS buffer, transferred to FACS tubes. Cells were stained with 2.5 μL DAPI (Biolegend, 422801), thoroughly mixed, and sorted for DAPI-/CD4+/TCRβ+ populations, followed by culture.

### Hybridoma activation

Splenic dendritic cells were used as antigen-presenting cells (APCs). C57BL/6 mice were injected intraperitoneally with 5×10^6^ FLT3L-expressing B16 melanoma cells to drive APC proliferation as previously described. Splenocytes were prepared 10-14 days after injection, and positively enriched for CD11c+ cells using Mouse CD11c Positive Selection Kit (STEMCELL, 18780). 2×10^4^ hybridoma cells were incubated with 10^5^ APCs and antigens in round bottom 96-well plates for two days. Cells were centrifuged at 800g for 5 min at 4°C and the pellet was resuspended in 50μL antibody mixture (0.2μL PE-Thy1.1(OX-7, Biolegend, 202524), 0.2μL APC-TCRβ in MACS buffer) and incubated for 15 min at 4°C. Cells were washed with 200μL PBS and centrifuged at 800 rpm for 5 min at 4°C. The pellet was resuspended in 200μL MACS buffer, transferred to FACS tubes. Cells were stained with 1μL DAPI (200×), thoroughly mixed. GFP induction in the DAPI-/Thy1.1+/ TCRβ+ hybridomas was analysed by flow cytometry as an indicator of TCR activation.

### Generation of TCR transgenic mice

Cb4-7 TCRα and TCRβ sequences of were cloned into the pTα and pTβ vectors provided by D. Mathis respectively. TCR transgenic mice were generated by the the animal facility at USTC. Positive pups were genotyped by testing TCR Vβ8.1/8.2 expression on T cells from peripheral blood.

### Synthetic peptides

Synthetic peptides representing antigens presented by IECs for hybridoma activation assays were synthesized by and purchased from Top-peptide (Shanghai). All peptides were above or equal to 95% purity (HPLC).

### Tetramer staining

∼1×10^6^ sorted T cells were stained by PE and APC-conjugated Cb_98-112_: I-A^b^ tetramers at 37°C for 2 hours in the dark. Then cells were washed twice and enriched by using anti-PE beads (Miltenyi Biotec) and anti-APC beads (Miltenyi Biotec). The enriched T cells were stained with Zombie Aqua™ dye as described above and analyzed by flow cytometry.

### Cell transfer

Single cells were collected from pooled spleens and lymph nodes of 8-10 week CD45.1 Cb4-7 Tg IL-10^GFP^/FOXP3^RFP^ dual reporter mice. Naïve CD4^+^ T cells were isolated by negative magnetic selection (Biolegend) according to manufacturer’s protocols. Isolated naïve CD4^+^ T cells were transferred into CD45.2 IL-10^GFP^/FOXP3^RFP^ dual reporter mice or *H2-Ab1^ΔIEC^, H2-Ab1^ΔDC^* and *H2-Ab1^ΔB^* mice crossed with IL-10^GFP^/FOXP3^RFP^ dual reporter mice by intravenous injection (1×10^6^ per mice).

### Statistical analysis

Data analysis was processed and represented by GraphPad Prism 9. Statistical significance was determined by two-sided Student’s t-test, one-way ANOVA Dunnett’s multiple comparisons test, one-way ANOVA Tukey’s multiple comparisons test and two-way ANOVA Šídák’s multiple comparisons, as indicated in figure legends. A *P* value of less than 0.05 (confidence interval of 95%) was considered significant. NS, no significance; \**P* < 0.05 \*\**P* < 0.01, \*\*\**P* < 0.001 and \*\*\*\**P* < 0.0001. The sample sizes are stated in the figure legends to indicate biologically independent replicates used for statistical analysis.

### Data availability

The single cell RNA and TCR sequencing data and the bulk RNA-seq data will be available in the Gene Expression Omnibus (GEO) repository at the National Center for Biotechnology Information after publish. All other data supporting the findings of this study are available from the corresponding authors upon reasonable request.

## Extended Data Figure 1-7

**Extended Data Fig. 1.**
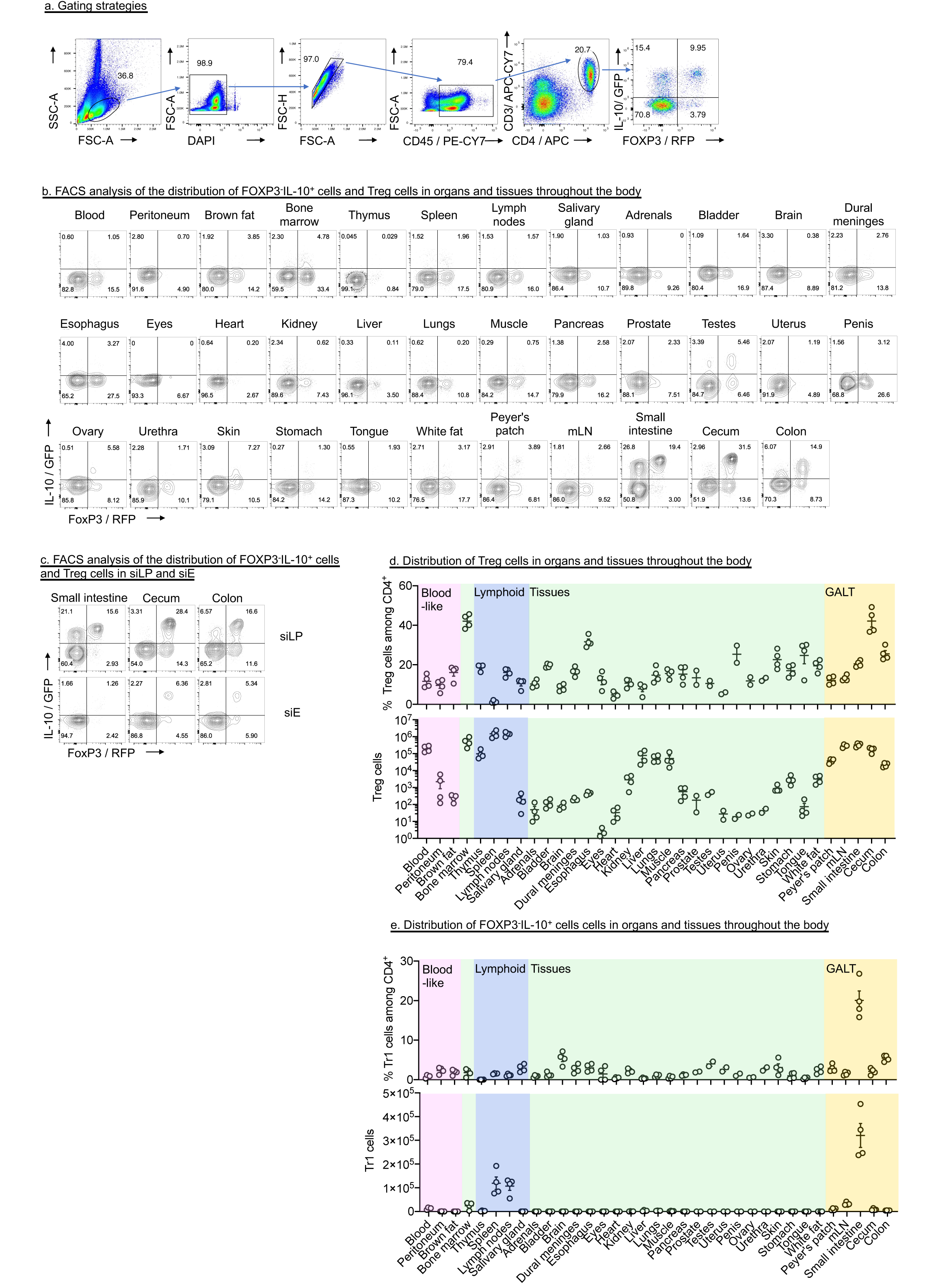
The distribution of Tr1 cells and Treg cells in various tissues and organs of mice. a, Gating strategy for CD4+ T cells isolated from the siLP of IL-10^GFP^/FOXP3^RFP^ dual reporter mice. b-d, Representative FACS plots (b and c) and quantification (d and e) of FOXP3 and IL-10 expression on gated CD45^+^ CD3^+^ CD4^+^ T cells isolated from the indicated tissue. Data are mean ± s.e.m and were pooled from two independent experiments.

**Extended Data Fig. 2.**
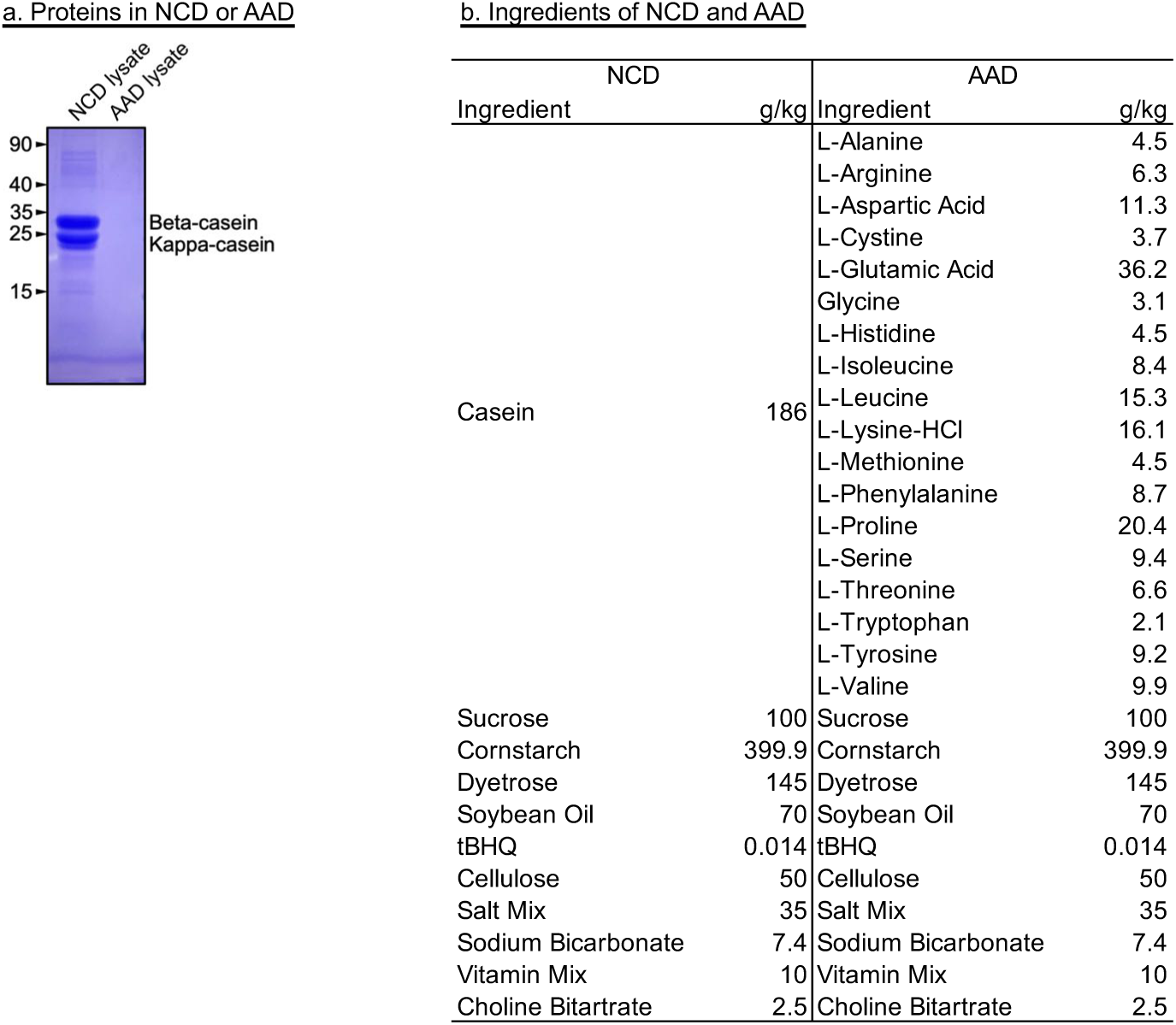
Diets used in the study. a, Coomassie blue staining indicated the protein composition of different foods. b, ingredients of NCD and AAD

**Extended Data Fig. 3.**
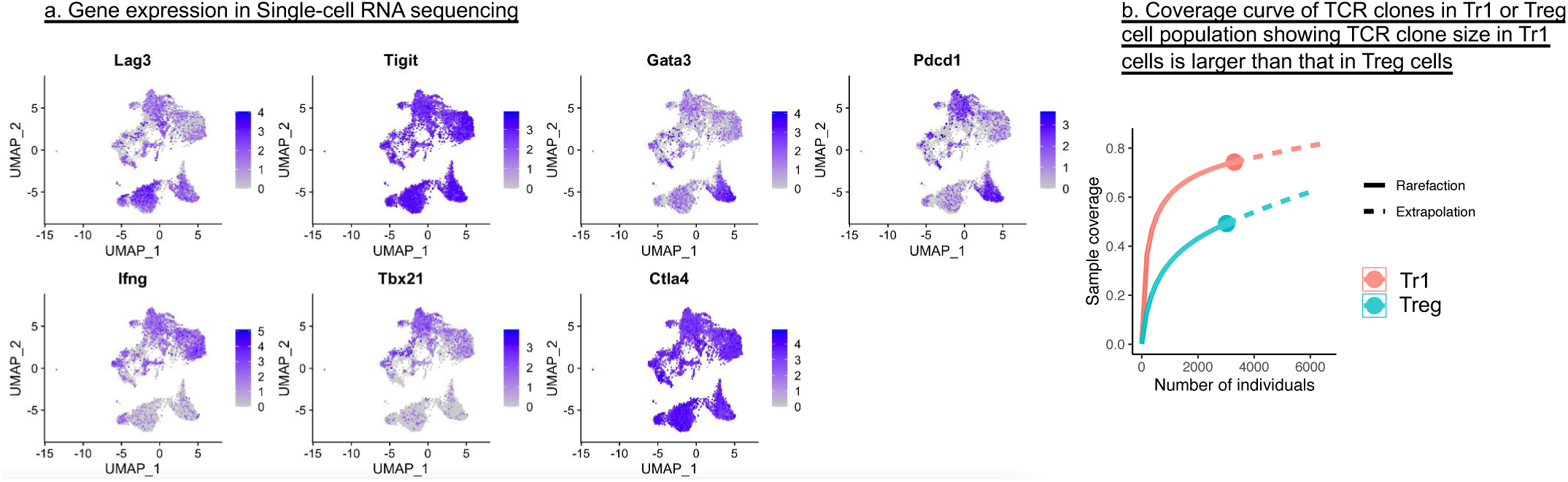
Single-cell RNA sequencing reveals that Tr1 cells and Treg cells have distinct transcriptional signatures and antigen specificity. a, Single cell expression of the indicated genes projected on the UMAP plot. b, Coverage curve of TCR clones in FOXP3^-^IL10^+^ Tr1 or FOXP3^+^IL-10^+^ Treg cell population isolated from the siLP.

**Extended Data Fig. 4.**
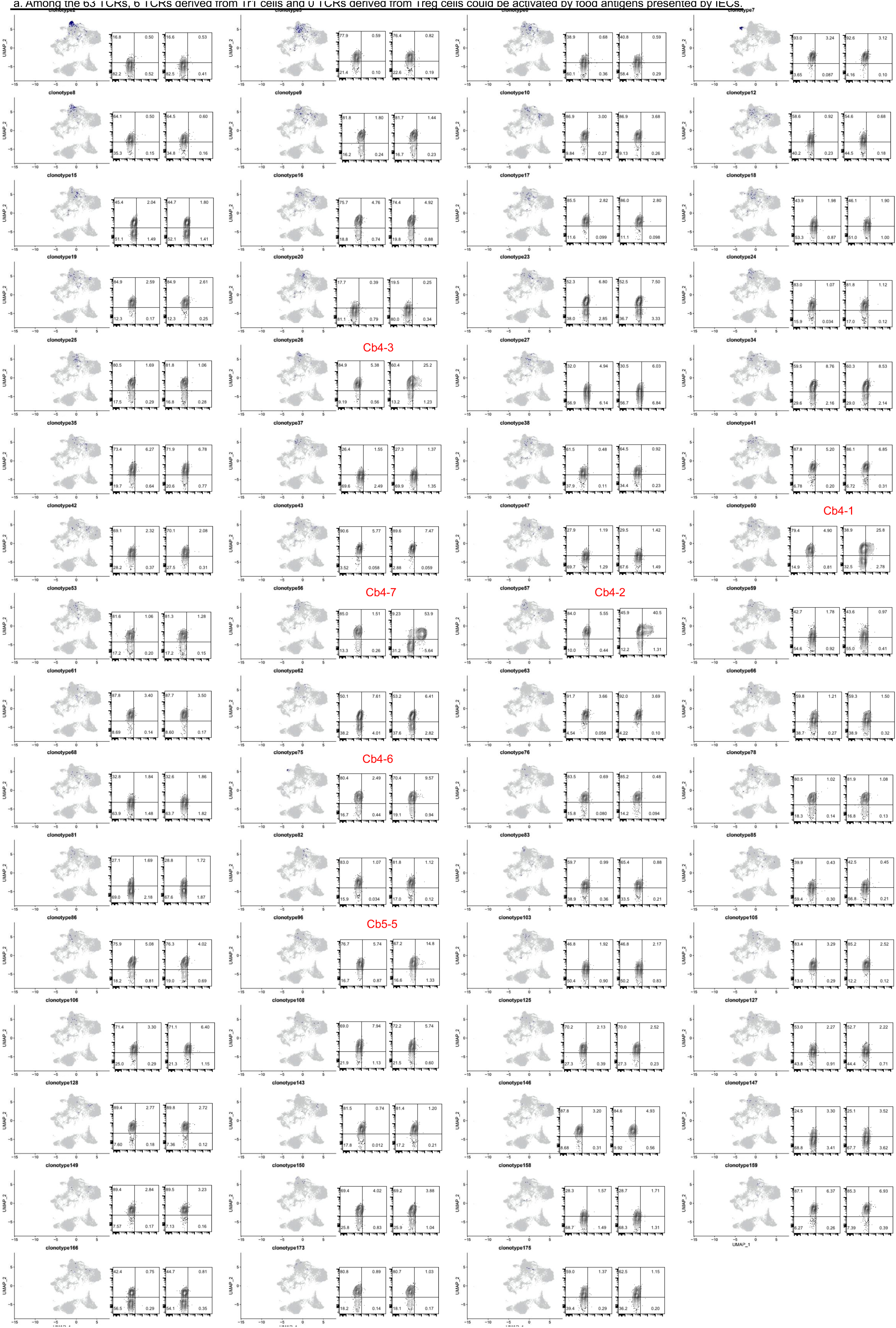
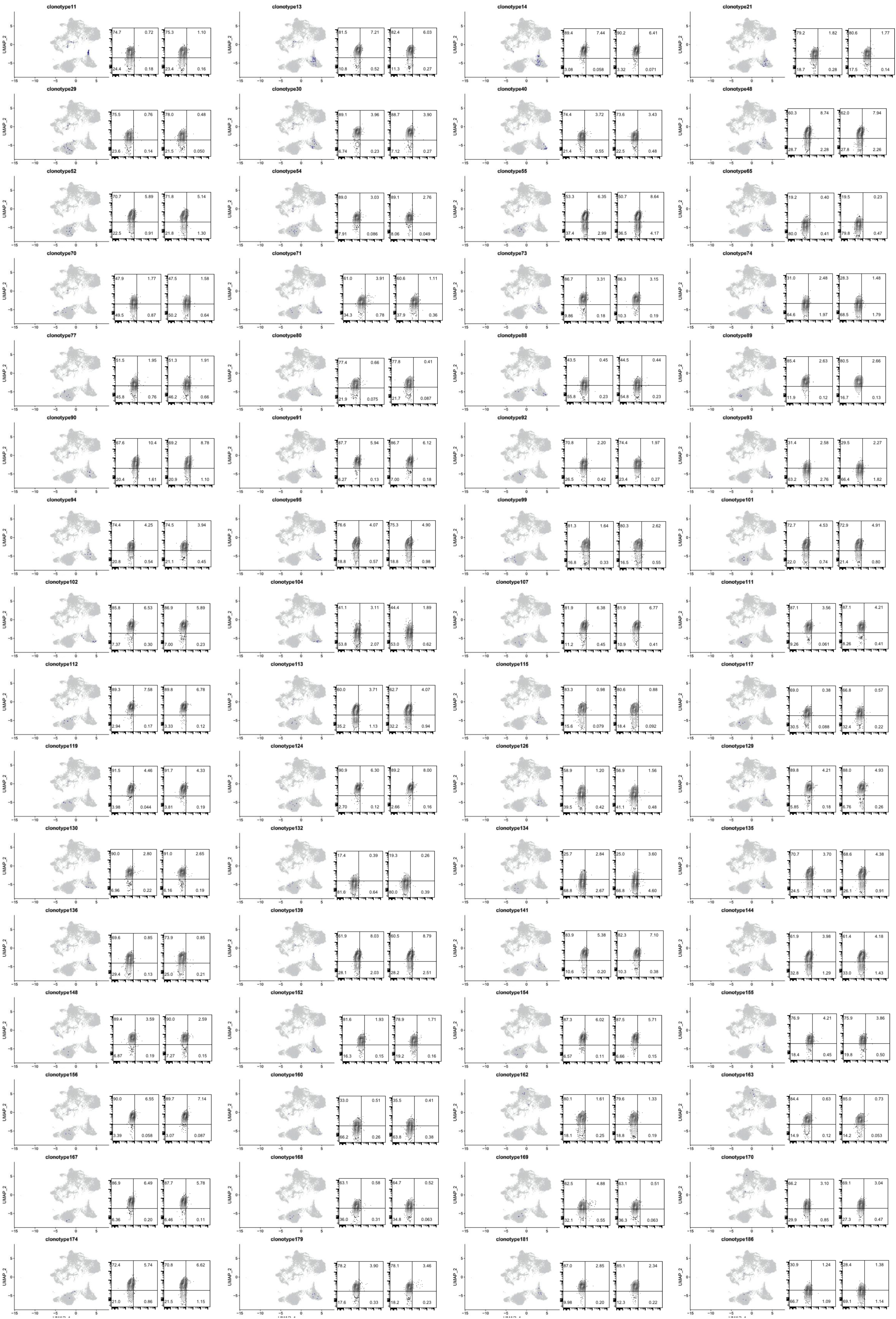
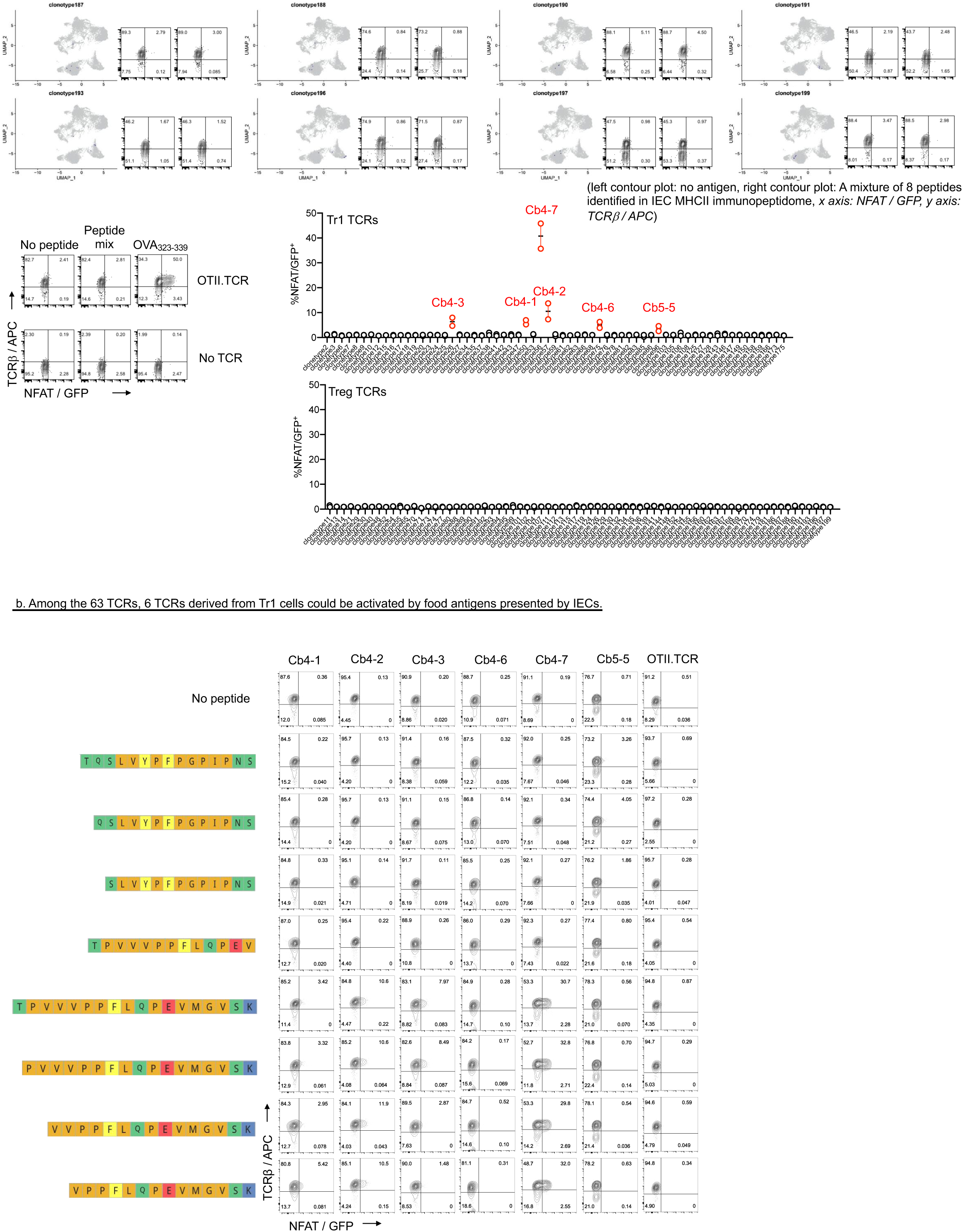
6 TCRs from the Tr1 TCR repertoire respond to dietary antigen presented by IECs. a, Stimulation of Tr1 or Treg TCR hybridomas by the mix of 8 identified IEC-presented food antigens. left: indicated T cell clone distribution projected onto the UMAP plot. right: representative FACS plots and quantification showing NFAT/GFP expression, which indicates the strength of TCR activation. The OTII TCR hybridoma served as a positive control, and the hybridoma without TCR transfection served as a negative control. 6 TCRs from the Tr1 TCR repertoires can be activated by the peptide mixture. b, Stimulation of 6 Tr1 hybridomas that respond to the peptide mixture by the individual peptide.

**Extended Data Fig. 5.**
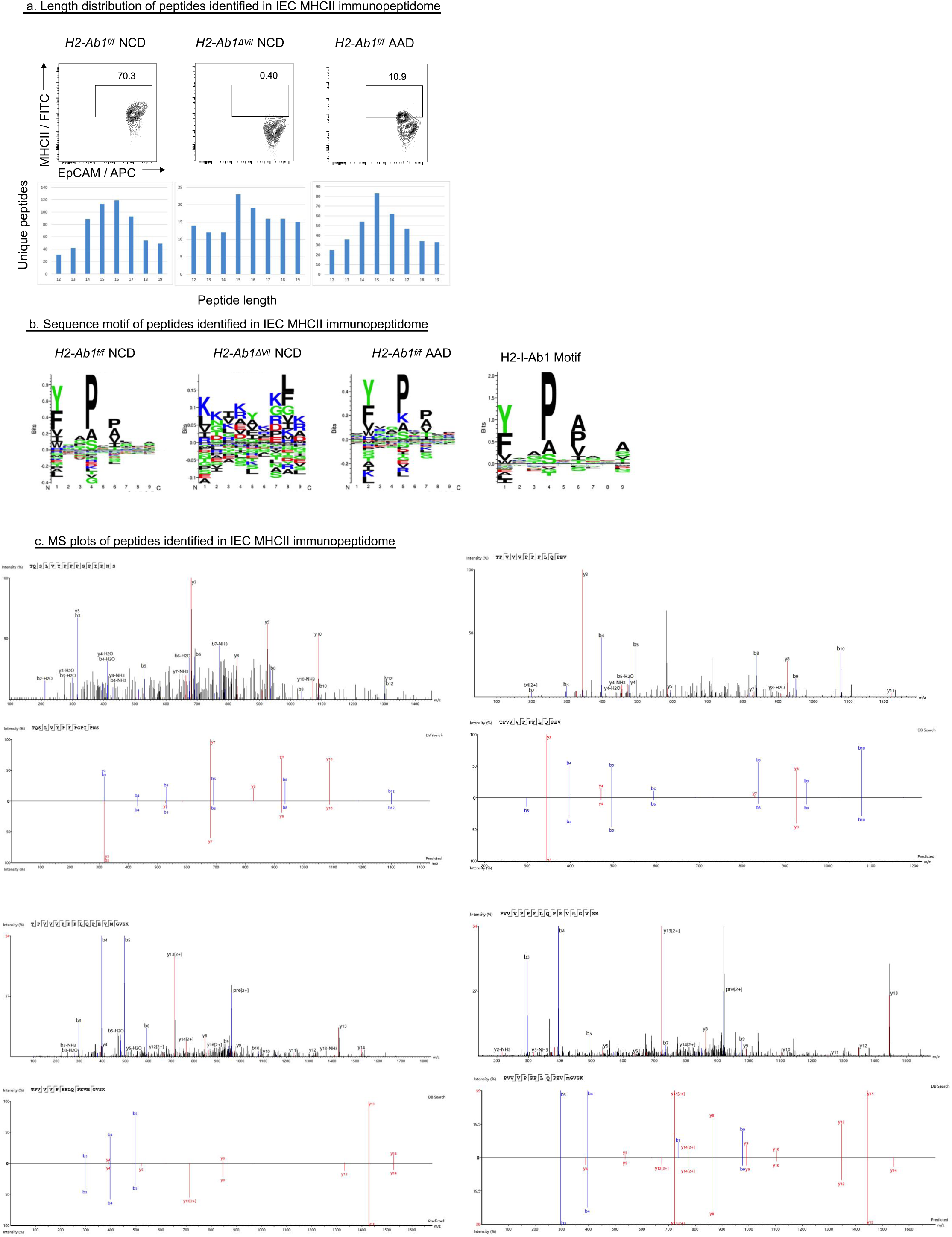
IECs present dietary antigens. a, FACS plots showing MHCII expression on IECs isolated from the indicated mice (upper panel). Length distribution of unique peptides identified in the indicated MHCII peptidome sample. b, Sequence motif of peptides identified in IEC MHCII immunopeptidome. c, Tandem mass spectra of the food peptides identified in the IEC-MHCII peptidome.

**Extended Data Fig. 6.**
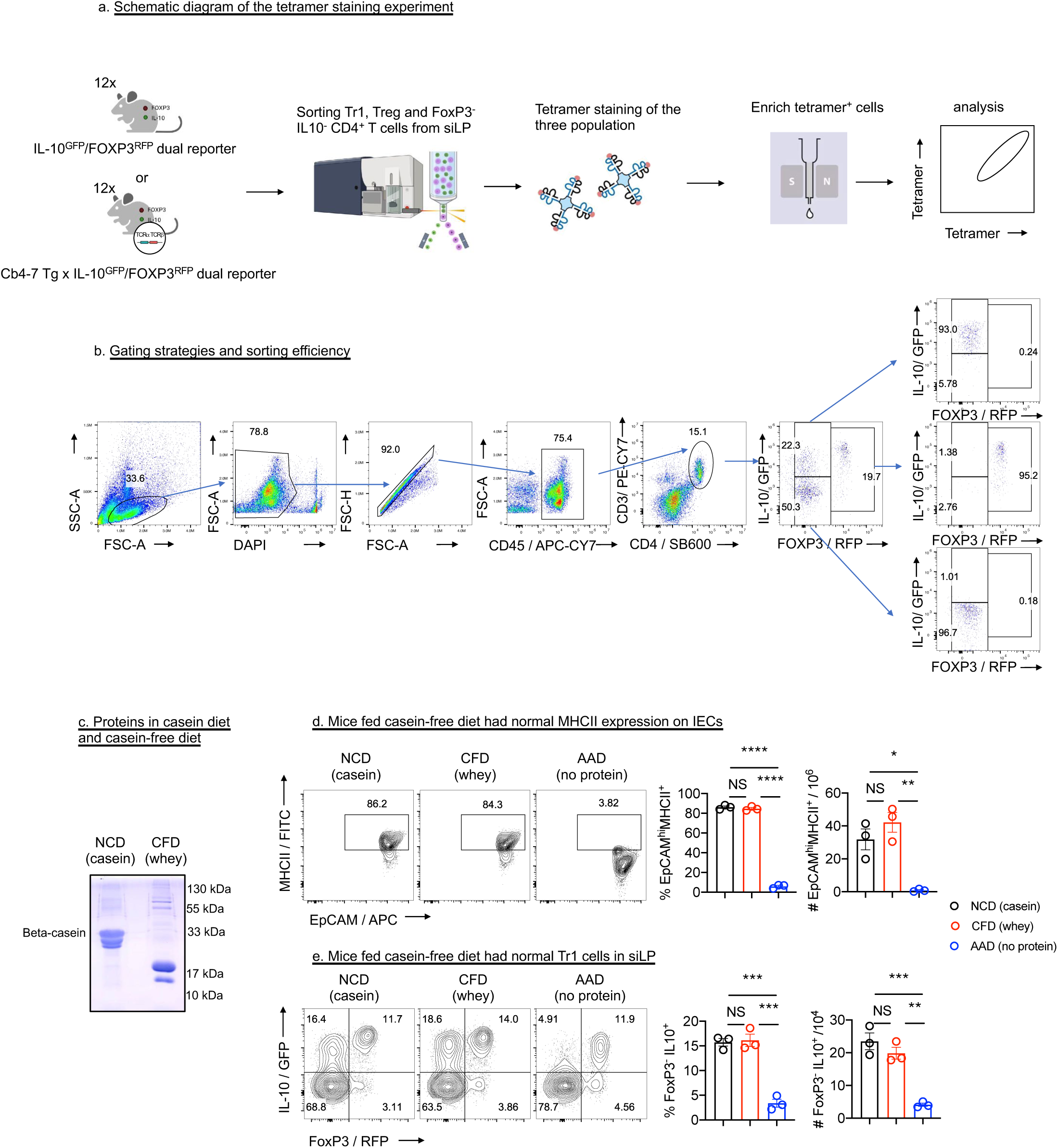
Tetramer staining of casein-specific T cells. a, Schematic diagram of the tetramer staining experiment. b, FACS plot showing the sorting efficiency of FOXP3^-^IL10^+^ Tr1 cells, FOXP3^+^ Treg cells and FOXP3^-^IL10^-^ T cells that isolated from the siLP of IL-10^GFP^/FOXP3^RFP^ dual reporter mice. c, Coomassie blue staining indicated the protein composition of NCD and CFD. d, Representative FACS plots and quantification of MHCII expression on IECs isolated from mice fed the indicated diets (n=3). e, Representative FACS plots and quantification of FOXP3 and IL-10 expression on gated CD45^+^ CD3^+^ CD4^+^ T cells isolated from the siLP of mice fed the indicated diets (n=3). Data are representative of three independent experiments. Data are mean ± s.e.m and were analyzed by one-way ANOVA Tukey’s multiple comparisons. NS, no significance; \**P* < 0.05, \*\**P <* 0.01, \*\*\**P* < 0.001, \*\*\*\**P* < 0.0001.

**Extended Data Fig. 7.**
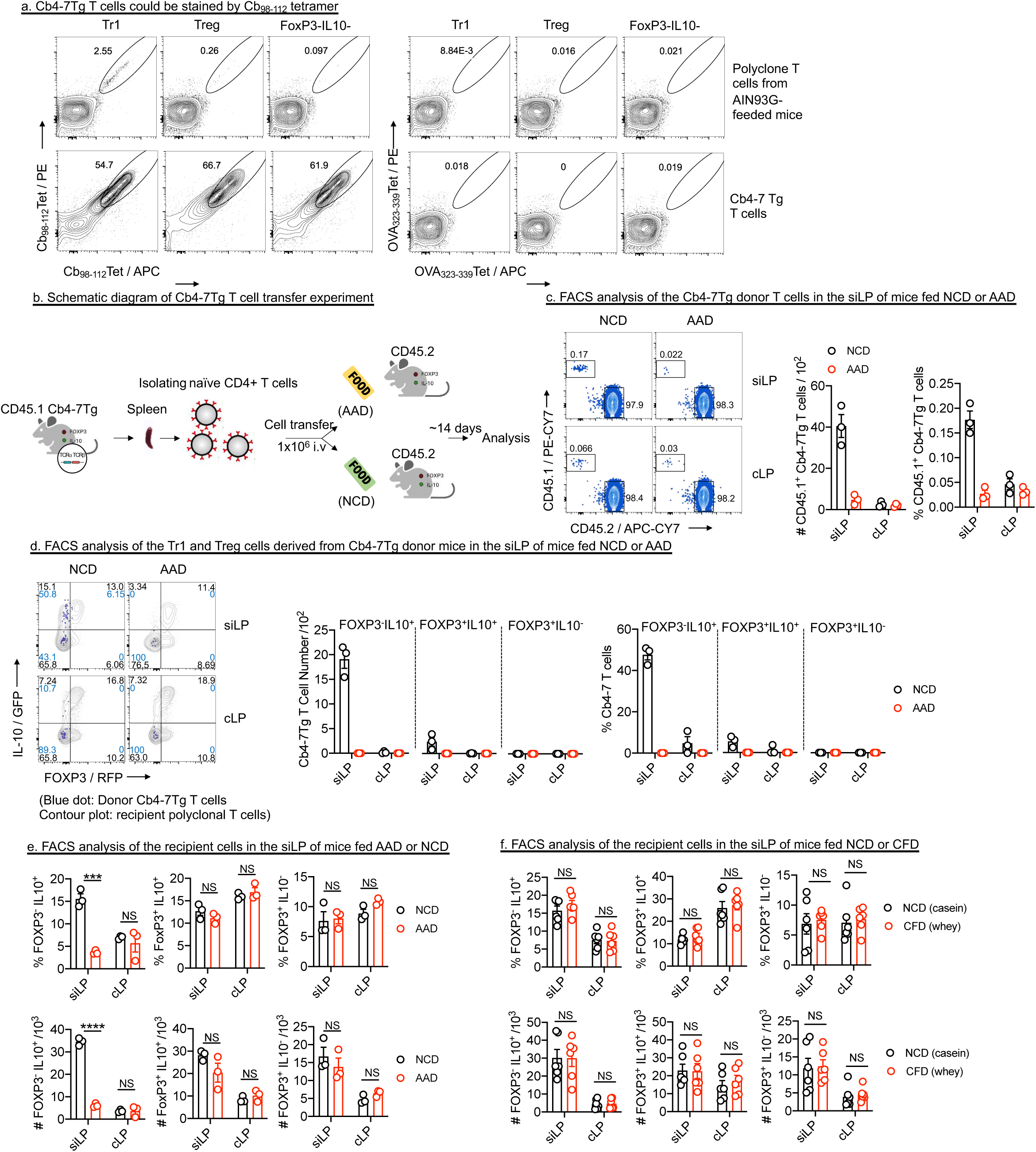
Food-specific T cells developed into Tr1 cells in siLP. a, Representative FACS plots and quantification of tetramer-positive Tr1 cells, Treg cells and FOXP3^-^IL-10^-^ cells sorted from the siLP of the indicated mice. b-d, CD45.1^+^ Cb4-7 Tg T cells were transferred into CD45.2^+^ hosts fed NCD or AAD. (b) Experimental schematic. (c) Representative FACS plots and quantification of donor CD45.1^+^ Cb4-7 Tg T cells in siLP or cLP of host mice. (d) Representative FACS plots and quantification of FOXP3 and IL-10 expression in donor (CD45.1^+^) and host polyclonal CD4^+^ T cells from siLP or cLP. e, quantification of FOXP3 and IL-10 expression on gated CD45.2^+^ CD3^+^ CD4^+^ host T cells isolated from the mice fed AAD or NCD. f, quantification of FOXP3 and IL-10 expression on gated CD45.2^+^ CD3^+^ CD4^+^ host T cells isolated from the mice fed NCD or CFD. Data are representative of at least three independent experiments. Data are mean ± s.e.m and were analyzed by two-way ANOVA Šídák’s multiple comparisons test, NS, no significance; \*\*\**P* < 0.001, \*\*\*\**P* < 0.0001.

